# Parallel clines in native and introduced ragweed populations are likely due to adaptation

**DOI:** 10.1101/773838

**Authors:** Brechann V. McGoey, Kathryn A. Hodgins, John R. Stinchcombe

**Author notes:** x.

## Abstract

As introduced species expand their ranges, they often encounter differences in climate which are often correlated with geography. For introduced species, encountering a geographically variable climate sometimes leads to the re-establishment of clines seen in the native range. However, clines can also be caused by neutral processes, and so it is important to gather additional evidence that population differentiation is the result of selection as opposed to non-adaptive processes. Here, we examine phenotypic and genetic differences in ragweed from the native (North America) and introduced (European) ranges. We used a common garden to assess phenotypic differentiation in size and flowering time in ragweed populations. We found significant parallel clines in flowering time in both North America and Europe. Height and branch number had significant clines in North America and, while not statistically significant, the patterns in Europe were the same. We used SNP data to assess population structure in both ranges and to compare phenotypic differentiation to neutral genetic variation. We failed to detect significant patterns of isolation by distance, geographic patterns in population structure, or correlations between the major axes of SNP variation and phenotypes or latitude of origin. We conclude that the clines seen for flowering time and size are most likely the result of adaptation.

## Introduction

Invasive species are both biological disasters and curiosities. In addition to the dual economic and ecological damage introduced plants can cause when they proliferate, they also offer opportunities to study evolutionary ecology during colonization (Callaway & Maron 2006). For example, invasions provide ideal systems to study the effects of reproductive isolation, how dispersal affects species distributions and the effect of a new individual species on an ecosystem (Sax *et al.* 2007). Evolutionary biologists have garnered insights from introduced species for decades by studying patterns of variation, interactions with native species, and comparing native and introduced populations (Huey *et al.* 2005). Clines in introduced populations offer an opportunity to study parallel evolution, the rate and predictability of local adaptation, and whether phenotypic divergence is due to selection or stochastic, non-adaptive forces (Huey *et al.* 2005; Samis *et al.* 2012; Colautti & Lau 2015). Here, we use a field common garden experiment and genotyping-by-sequencing (GBS) to investigate clines in introduced and native ragweed populations with the goal of distinguishing between adaptive and non-adaptive mechanisms underlying clinal variation.

Incorporating an evolutionary perspective into invasion biology is critical to understanding the course an invasion has taken and how it might continue to unfold. For example, the capacity of an introduced population to adapt to its new climate is important for its capacity to persist and spread (Colautti & Barrett 2013). Adaptation to climate variables often lead to geographic differentiation, as climate and geography are strongly correlated (Endler 1977). The common pattern of a gradient in traits or alleles over a geographic range (Huxley 1938) is often interpreted as evidence of adaptive differentiation. Clines can be found in both Mendelian and quantitative traits and there are hundreds of examples across a wide variety of taxa (Campitelli 2013). Clines in introduced species, especially those that mirror geographic variation in the native range, are often perceived as evidence that introduced populations have adapted to their new environments (Samis et al. 2012; Colautti and Barrett 2013). However, processes other than adaptation can also be responsible for both phenotypic and genetic clines, and need to be controlled for (Keller and Taylor 2008). For example, phenotypic clines observed *in situ* could be caused by plastic responses to environmental variables, especially in plant species (Huxley 1938), and neutral processes present through colonization could also produce clines (Vasemägi 2006; Keller *et al.* 2009; Santangelo *et al.* 2018).

Distinguishing between the possible forces underlying clines can be achieved in several ways. Plasticity can be excluded by using a common garden to ensure that any differences between populations have a genetic basis (Lucek *et al.* 2014). Linking clines with natural selection and demonstrating a correspondence between the direction of selection and variation in phenotypes can also strengthen the case that a cline is the result of adaptation (Etterson *et al.* 2008). Parallel clines can also be interpreted as evidence of adaptation (Samis et al. 2012). Neutral markers can be used to rule out drift and support hypotheses that differences are due to adaptation (Keller & Taylor 2008; Kooyers & Olsen 2012; Lima *et al.* 2012; Campitelli and Stinchcombe 2013; Le Gros *et al.* 2016). Likewise Q_ST_-F_ST_ comparisons can be used to compare molecular and quantitative genetic variation. Whereas F_ST_ examines differentiation at neutral markers, Q_ST_ is analogous but quantifies population divergence for quantitative traits (Spitze 1993). Meta-analyses have found that genetic differentiation among introduced populations are common and on average do not differ much in magnitude from divergence between native populations(Colautti and Lau 2015). By using these methods, researchers have bolstered the argument that rapid adaptation occurs in non-native species from plants to fruit flies (Huey *et al.* 2000; Montague *et al.* 2008; Samis *et al.* 2012; Colautti & Lau 2015).

*Ambrosia artemisiifolia* (ragweed) is a globally invasive species with a wide range in its native continent of North America and a presence in Europe, Asia and Australia (Friedman & Barrett 2008). Past work on *A. artemisiifolia* has demonstrated parallel clinal patterns in flowering time in the native and introduced ranges (Hodgins & Rieseberg 2011). In this experiment, we examine variation in several quantitative traits across geography. To determine how quantitative variation may have been impacted by neutral processes, we use single neutral polymorphism (SNP) data to assess neutral genetic variation. We ask the questions: **Are there clinal patterns in quantitative traits, in the introduced and native ranges? Are the patterns consistent with a history of selection in the introduced range, or non-adaptive processes?** Our results corroborate, using independent biological samples, experiments, and analytical approaches, past results (Hodgins and Rieseberg 2011; van Boheemen et al. 2018) of clinal variation in ragweed being likely due to adaptive differentiation rather than stochastic processes.

## Methods

### Study species

*Ambrosia artemisiifolia* (common ragweed) is an annual outcrosser in the Asteraceae family (Bassett & Crompton 1975; Friedman & Barrett 2008). Ragweed is thought to have originated in the plains of North America, and then spread eastward (Bassett and Crompton 1975). In the modern era, ragweed has been accidently introduced to Europe, Asia and Australia (MacKay & Kotanen 2008). In Europe, ragweed has been present since at least the mid-1800s, but propagule pressure (composite measure of the individuals or seeds released in an introduction and the number of introductions (Lockwood *et al.* 2005)) increased dramatically in the mid-20^th^ century when ragweed seeds contaminated grains shipped from the Americas to Europe (Chauvel *et al.* 2006). The geopolitical situation during that period meant that imports were coming from different areas into Western versus Eastern Europe. This resulted in two invasion centres, with distinct genetic origins (Gladieux *et al.* 2011). In France, the epicenter of the invasion is the Rhône valley, where ragweed grows in very large populations along riverbanks (Chauvel *et al.* 2006; Thibaudon *et al.* 2013). The most impacted nation is Hungary, and in Eastern Europe ragweed populations now extend north up into Poland and south into the Baltic states (Prank *et al.* 2013). Ragweed is considered one of the most problematic invaders in Europe: it causes allergies, and is a significant agricultural weed. In Hungary, it is the most widespread weed in surveys (Kiss & Béres 2006) and over 80% of arable land is affected (Buttenschøn *et al.* 2010).

### Seed collection and preparation

In the autumns of 2012 and 2013, we collected seeds from a total of 20 native and 18 introduced populations (Figure 1; population coordinates in Appendix Tables A1 and A2). In both ranges the populations spanned ∼7.5 degrees in latitude. These populations ranged from small (5 individuals) up to tens of thousands of individuals. When populations had fewer than twenty individuals, we collected seeds from all the plants. For larger populations, we collected from a random subset. Using methods adapted from Willemsen (1975) and Jannice Friedman (Queen’s University, pers. comm.), we stratified seeds at 4°C for six months in plastic bags filled with silica and distilled water.

**Figure 1.**
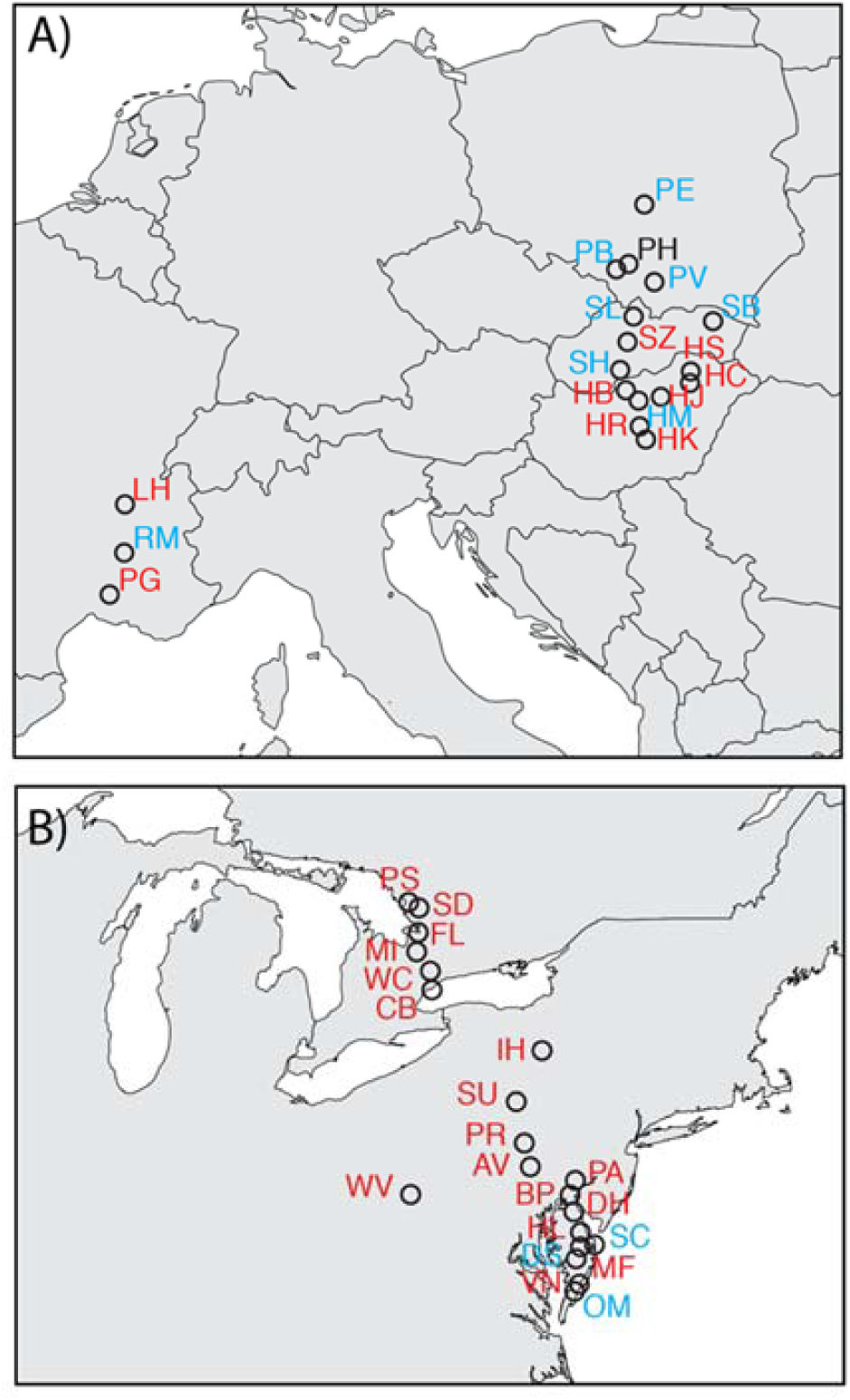
Map of population *Ambrosia artemisiifolia collection* sites from. (A) Native North American range (B) Invasive European range. Collection sites used only for the common garden are shown in blue. Sites used for both the common garden and SNP study are shown in red.

### Common garden

For this experiment we randomly chose 10 maternal families per population, except in cases where there were fewer than 10 families, in which case we used all available. We chose three germinants from each family (one for each of three blocks) for a total of 1110 plants. Each individual was randomly assigned a position within a block. We planted germinants in seedling trays and kept in a greenhouse where we sprayed and bottom watered them for three weeks. At the end of June 2014, we planted seedlings into a plowed field at the Koffler Scientific Reserve (www.ksr.utoronto.ca; 44.803°N, 79.829°W). Blocks were subdivided into plots, each containing 64 plants in an 8*8 configuration, except for the final plot. We continued to remove interspecific vegetation and provide the seedlings with water for four weeks after transplantation to promote establishment.

### Phenotypic traits

We measured final height, final number of branches, and date of first flower. Since the vast majority of ragweed plants are monoecious (Bassett and Crompton 1975), we measured proxies of both male and female fitness. Male reproductive effort was estimated as the total inflorescence length, which is correlated with pollen production (Fumanal *et al.* 2007). We estimated female reproductive output, using seed mass, which is highly correlated with seed number (r^2^=0.96, p<0.001) (MacDonald and Kotanen 2010). We ran initial models including latitude and block to assess if block had a significant effect on traits; block was not significant for any trait, and was thus dropped in subsequent models. To test for clines in phenotypic traits, we used linear regressions. For each continent, we conducted regressions for three phenotypic traits (height, flowering time and branch number) and the two fitness traits on latitude. Similar results were obtained using population means for traits and latitude. Unless otherwise specified below, statistical analyses were conducted in R (R Development Core Team 2016).

### Neutral markers

We collected leaf material from 180 ragweed plants from 26 populations (9 introduced, 17 native) grown in seedling trays in growth chambers at the University of Toronto. These plants were from a subset of the populations used in the common garden, but were separate plants. GBS library prep was conducted by the Hodgins lab at Monash University Australia. In brief, DNA was digested from the dried leaves and adapters were ligated to the strands. A double enzyme digest with Pst1 and Msp1 was implementing using the same protocol as in van Boheemen et al (2018). DNA libraries were sent to Genome Quebec for sequencing on an Illumina HiSeq 2500, using PE125 sequencing.

We used Stacks (Catchen *et al.* 2013) and Bowtie 2 (Langmead & Salzberg 2012) to demultiplex, align to a reference genome provided by the Hodgins lab and to calculate population genetic metrics. We checked the sequence quality using FastQC (Andrews 2010) and samtools (Li *et al.* 2009).We converted between multiple formats using PGD Spider (Lischer & Excoffier 2011), samtools (Li et al. 2009), Bowtie 2 (Langmead and Salzberg 2012), admixr (Petr n.d.) and custom python and bash scripts (see Github). To prepare for STRUCTURE and isolation by distance (IBD) analysis we filtered snps using vcf tools (Danecek *et al.* 2011). We excluded snps with a minor allele count lower than 4 (equivalent to 2.2%) or with data missing in greater than 20% of samples. These thresholds were average to conservative based on those used in previous studies (McGrath 2014; Huang *et al.* 2014; Taylor *et al.* 2014; Sawler *et al.* 2015; Beck & Semple 2015; Ilut *et al.* 2015; Martin *et al.* 2016; Mondon *et al.* 2017).We set the significance threshold for Hardy-Weinberg equilibrium at 1e-5, which was the midpoint used in a review of past studies (Anderson *et al.* 2010). Since very rare alleles could still be important for assessing population structure (Linck & Battey 2017), we evaluated whether inclusion of them altered the results, and found that they did not. We present results with the exclusion of SNPs with rare minor allele counts <2.2%.

### Population Structure and Geography

To conduct a STRUCTURE analysis (Pritchard *et al.* 2000) while taking advantage of parallelization, we used StrAuto (Chhatre & Emerson 2017).We conducted separate analyses for the two continents, with five replicates at K=1-6 for each. In addition, we ran a STRUCTURE analysis using data from all the populations together. To visualize the STRUCTURE output we used the default settings of the web-based program Cluster Markov Packager Across K (CLUMPAK) (Kopelman *et al.* 2015) and the R package *pophelper* (Francis 2017).

To examine isolation by distance (IBD), we used the R package *adegenet* to test for IBD in each continent separately (Jombart 2008). *Adegenet* uses a Mantel test between matrices of genetic and geographic distances to assess if more spatially disparate populations are also more genetically divergent.

In addition to STRUCTURE and IBD, patterns in genetic data can also be understood with principal component analysis (Cavalli-Sforza *et al.* 1994; Patterson *et al.* 2006; McVean 2009; Josephs *et al.* 2018). We performed PCA on SNP data from the native and introduced ranges to test for correlations between major axes of SNP variation and the phenotypic traits of interest, and latitude. To extract principal components from SNPs we used the R package LEA (Frichot & François 2015). The vcf files were converted to geno files using the vcf2geno function. We then ran PCAs of all available SNPs separately for the two continents. To determine which principal components should be retained in subsequent analyses, we used Tracy-Widom tests (Patterson *et al.* 2006). For each population, we calculated a PC score along each significant axis of SNP variation, and then used these PC scores to test for associations with traits or geography. Specifically, to determine whether axes of neutral SNP variation were related to either geography or phenotype with linear regressions. For each continent, we regressed each significant principal component on latitude and the three phenotypic traits.

### Descriptive Population Genetic Statistics

To explore population genetics of the native and introduced ragweed populations, we used the programs Genodive and Splitstree (Hudson 1998; Meirmans & Van Tienderen 2004). We converted vcf files for each continent, and for the entire dataset, to the genetix format and then imported them into Genodive. We then used Genodive to estimate F_ST_, population genetic summary statistics (including observed and expected heterozygosity and fixation indices), and to run an AMOVA to partition genetic variation between the range, population and individual levels. We used Splitstree to construct a neighbornet for all the individuals (Hudson 1998).

### Data availability

Upon acceptance, accession numbers for sequences deposited in the short read archive will be placed here (https://www.ncbi.nlm.nih.gov/sra). A summary of our bioinformatics pipeline, including code and the original fastq files is available here (https://github.com/brechann-mcgoey/ragweedGBS)

## Results

### Phenotypic traits

Plants from more southern latitudes flowered later and grew larger both in total height and branch number (see Figure 2). These clines were all significant for the North American populations, while only flowering time had a statistically significant association with latitude in Europe (see Table 1). There was only one significant cline for fitness traits, with more southern North American plants producing more fruits than those in more northern populations (Figure 3). Since more southern plants were also larger, this correlation may be driven by a relationship between size and latitude.

**Table 1.** Result of regressions of *Ambrosia artemisiifolia* traits on latitude in the invasive (European) and native (North American) ranges. *** Indicates P<0.01.

|  | Flowering time | Height | Total branches | Male fitness | Female fitness |
| --- | --- | --- | --- | --- | --- |
| **EUROPE** |  |  |  |  |  |
| **Estimate** | -1.545*** | -0.672 | -0.120 | -9.523 | -0.578 |
| **Std. error** | (0.319) | (0.578) | (0.180) | (7.236) | (0.387) |
| **Degrees of freedom** | 231 | 228 | 227 | 228 | 194 |
| **NORTH AMERICA** |  |  |  |  |  |
| **Estimate** | -4.694*** | -5.570*** | -0.333*** | -10.557 | -2.243*** |
| **Std. error** | (0.262) | (0.495) | (0.129) | (6.776) | (0.339) |
| **Degrees of freedom** | 287 | 285 | 284 | 285 | 262 |

**Figure 2.**
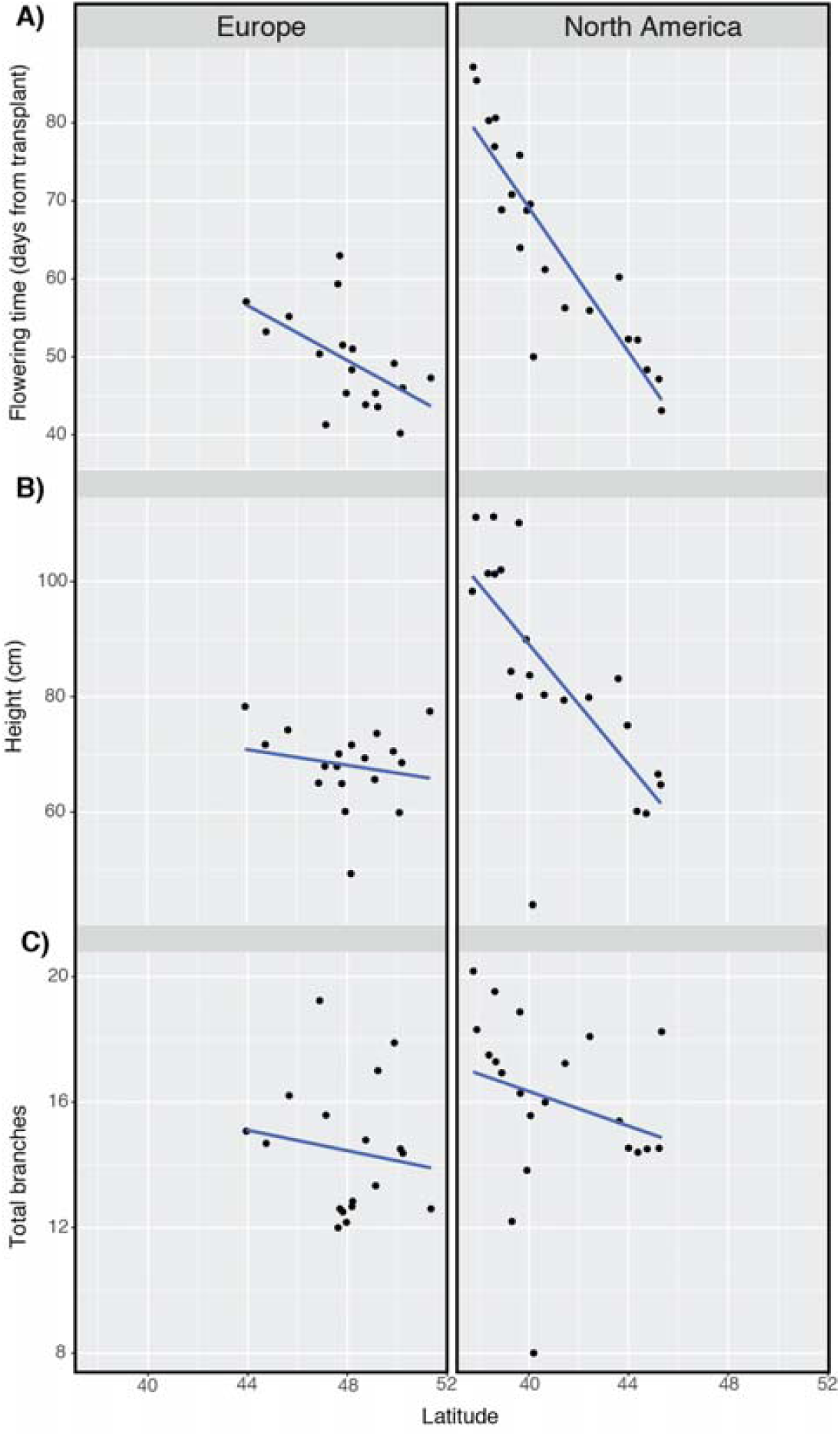
Phenotypic results for ragweed plants grown in common garden. (A) Flowering time (B) Height (C) Total branches on the y axes, the latitudes of origin on the x-axis.

**Figure 3.**
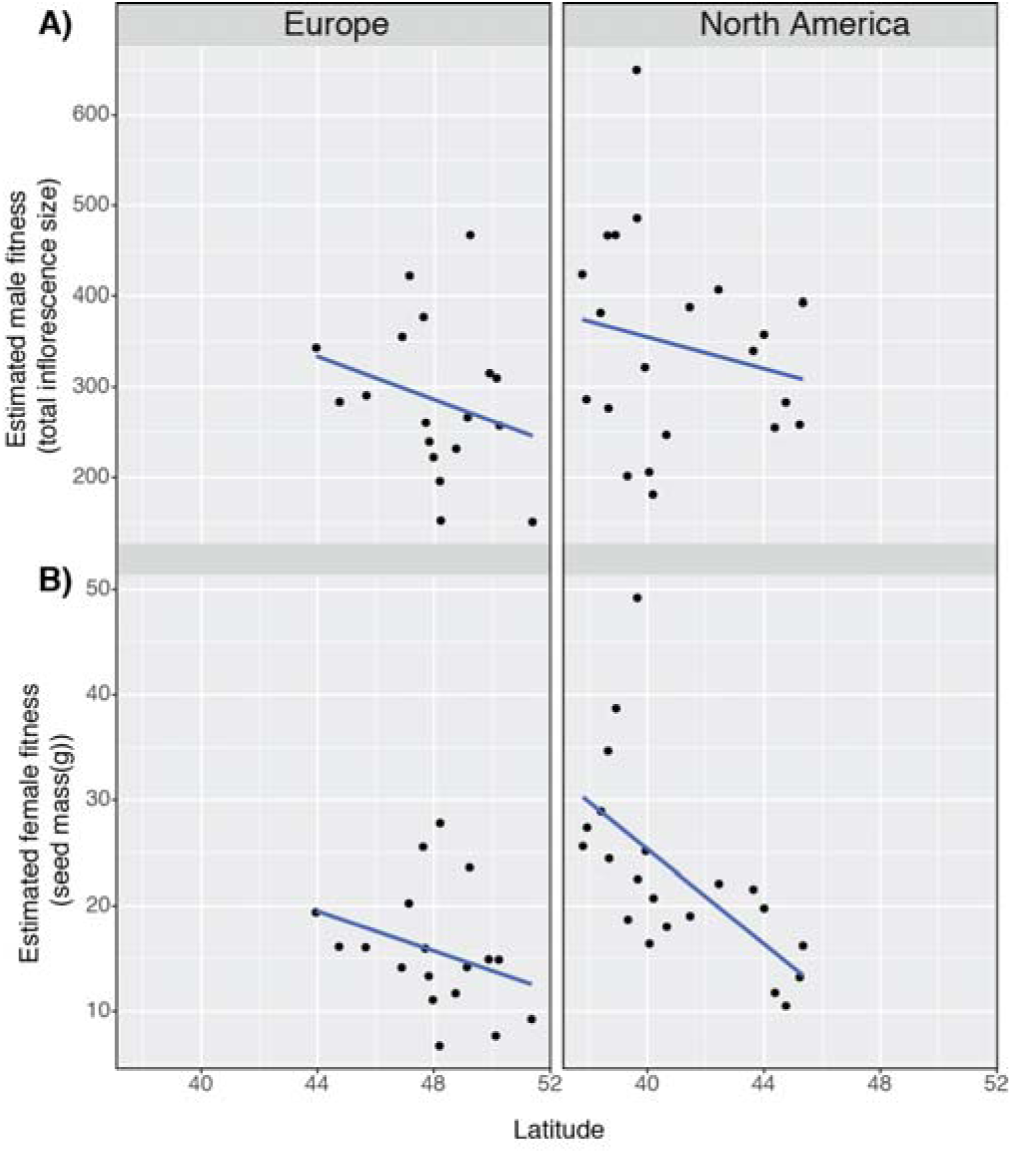
Female and male reproductive output results for ragweed plants grown in common garden. (A) Estimated male fitness (B) Estimated female fitness.

### Neutral markers

The Illumina analysis resulted in 250 033 952 total sequences which all passed a FastQC quality check. We started with 258 540 SNPs across all the populations. After filtering, we had 20 843 sites for Europe and 28 643 for North America. Our STRUCTURE analysis indicated that the European samples clustered in three groups, while the North American lines clustered in five groups. In Europe, there was no obvious geographic pattern to ancestry (see Figure 4A). In North America, two populations seem distinct, but otherwise there is not much geographic structure (see Figure 4B). There was no significant isolation by distance in either continent (p-value of 0.84 for Europe and 0.342 for North America). Our Splitstree analysis was consistent with the above results, showing populations were not clustered into distinct subgroups (Appendix Figure A1). The global STRUCTURE analysis for all populations found the most support for k=5. As with the by-continent analyses, little of the ancestry was geographically structured (see Appendix Figure A2).

**Figure 4.**
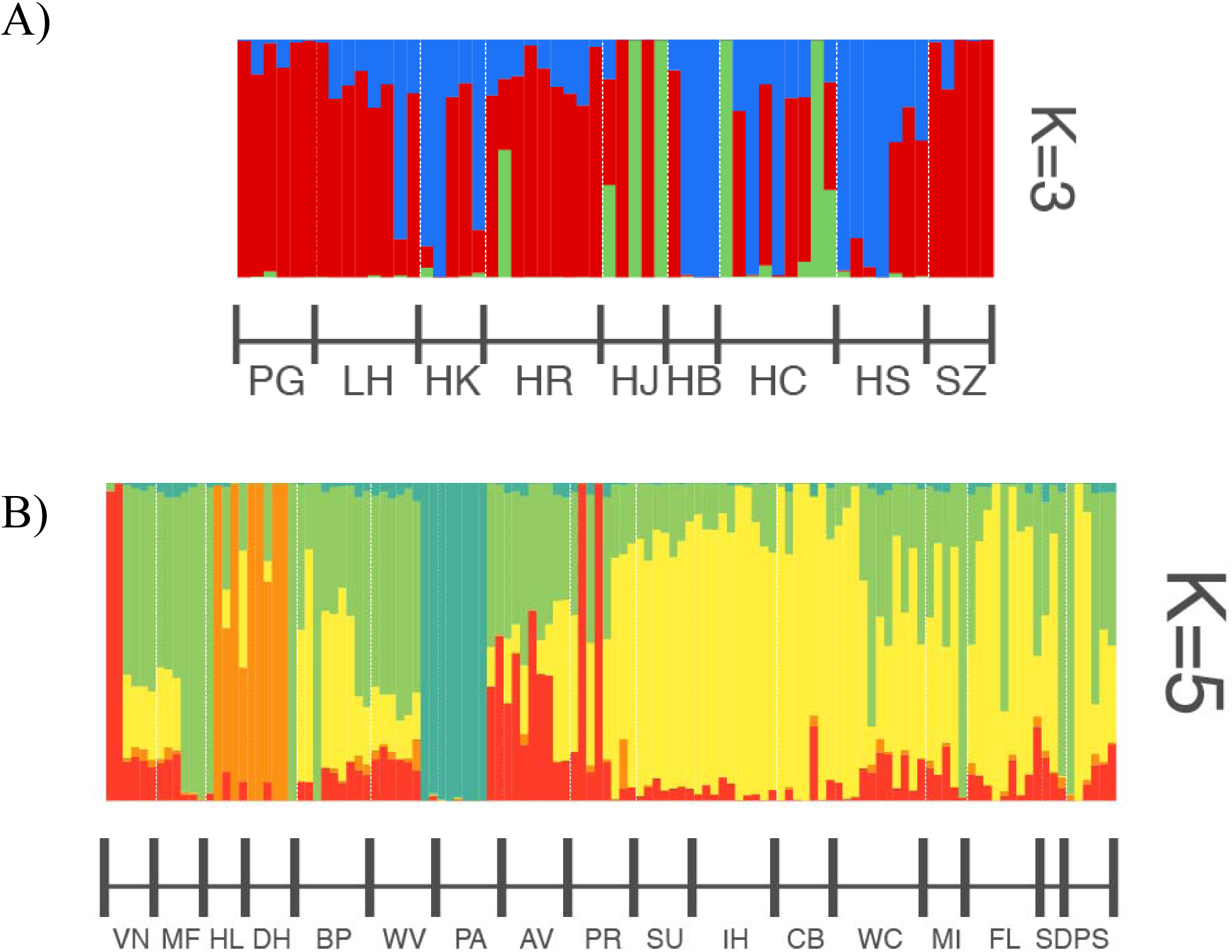
STRUCTURE results for ragweed populations (A) Europe (B) North America. Populations are ordered from lowest to highest latitudes (south to north).

Tracy-Widom tests indicated that there was one significant SNP principal component in Europe and seven for the North American dataset (North America: TW Statistic ≥ 1.029, P ≤ 0.04671 for PCs 1-7; Europe: TW Statistic = 6.255, P = 1.013E-06 for PC1). The significant European principal component explained 4.5% of genetic variation. Altogether, the significant North America principal components explained 14.6% of genetic variation. There were no significant relationships between significant SNP principal component scores and either latitude or phenotype (all p values >0.05). These results suggest that axes of (neutral) SNP variation are not related to either latitude, or the phenotypic traits, suggesting that latitudinal clines are unlikely to be generated by neutral or stochastic processes.

In keeping with the lack of geographic structure revealed by the Splitstree and Structure results, F_ST_ values were universally low (mean of 0.0639 (range of 0.003 to 0.213) for North America and 0.0566 (range of 0.018 to 0.12) for Europe, see Appendix tables A13& A4 for population specific results). Other population level statistics were consistent with the conclusion that there is low population differentiation in both ranges (see Appendix tables A5 & A6). An AMOVA of the global dataset indicated that very little variance was at the between continent or between population levels (see Appendix table A7). Total heterozygosity was lower in native (0.150) as compared to the invasive range (0.202). The expected heterozygosity within populations was also higher in the invasive range (0.192 vs 0.141 for the native range).

## Discussion

Introduced species offer a unique opportunity to address important questions in evolutionary biology (Sax et al. 2007; Yoshida et al. 2007). Adaptation is an important and sometimes overlooked aspect of invasions (Huey et al. 2005; Barrett 2015). New species introductions offer an excellent avenue to explore the rate and predictability of adaptation, a topic of great interest to evolutionary biologists (Huey et al. 2005). The reemergence of genetic clines in introduced ranges represents evidence that adaptation may occur quickly and result in predictable phenotypic differentiation (Leger & Rice 2007). Adaptation to local habitats and population differentiation can be critical to the ability of an invasive species to expand its range (Colautti and Barrett 2013). Here we detected phenotypic differentiation among native and introduced populations of common ragweed, and evidence that these clines are the results of local adaptation as opposed to neutral processes. We also examined the population structure and genetic diversity of both native and invasive populations.

### Clines as evidence of local adaptation

Past studies of ragweed have found geographic patterns in phenotype, including parallel clines in the native and introduced ranges (Hodgins and Rieseberg 2011, van Boheeemen et al. 2019). With populations from different parts of both Europe and North America than those used in previous studies, we corroborated their conclusions that flowering time patterns have been reproduced in the introduced range. We present evidence that these patterns are almost certainly the results of selection as opposed to neutral processes. Unlike flowering time, patterns of SNP variation were not correlated with latitude, and principal components of genetic variation were not correlated with the phenotypic traits we examined, including flowering time. Overall there was little population structure and low F_ST_ values were identified in both ranges. Van Boheemen et al. (2019) detected repeated latitudinal divergence in a host of life history and size traits in Australian, North American, and European samples of ragweed, as well. In their case, they detected these clines when controlling for structure coefficients (so-called q-values) as a measure of neutral population genetic structure. Here we used independent biological samples, phenotyped plants in a field common garden in the native range (as opposed to growth chamber), and used alternative means of assessing neutral population genetic structure (PCA of the SNP matrix). That we found qualitatively similar results—clinal differentiation above and beyond what could be explained by neutral processes—all make it unlikely that the cline in flowering time is the results of drift.

One caveat of this study is that all plants were grown from seed and therefore subject to maternal effects. However, we think it unlikely that maternal effects could explain the phenotypic patterns presented here. The population level maternal effects would all need to be in a consistent direction across latitude. Maternal effects are more pervasive in early-life history stages (Rossiter 1996; Montague *et al.* 2008), while here we focused on traits at the end of the annual life cycle. It is also unlikely that maternal effects would be responsible for differentiation at such large geographic scales (Montague *et al.* 2008). In addition, the clines found here were consistent with those of our previous work (McGoey and Stinchcombe 2018), which included fewer populations but did remove the impact of maternal effects.

Ragweed has been present in Europe for ∼250 years (Chauvel *et al.* 2006). In as many generations, it has spread and proliferated, especially in France, Italy and the Pannonian Plain (Thibaudon *et al.* 2013). Our previous research demonstrated that invasive ragweed populations possess ample additive genetic variation, in spite of any bottlenecks and founder effects during the colonization process. Given the results shown here, it must also be concluded that this quantitative genetic variation has been preserved in the face of selection and adaptation as well.

Past studies of population differentiation, including one examining ragweed in its introduced range, have used Q_ST_-F_ST_ comparisons to assess geographic variation (Leinonen *et al.* 2008; Chun *et al.* 2009). As with our study, Chun and colleagues found the invasive ragweed populations had low F_ST_ values and geographic structure (Chun *et al.* 2009). Their Q_ST_-F_ST_ analysis indicated there was significant diversifying selection acting on ragweed in its introduced range. The conclusions of Chun and colleagues align with our own, that population differentiation in the invasive range is the result of adaptive evolution as opposed to neutral processes. Q_ST_-F_ST_ comparisons are very interesting but fraught with technical challenges and statistical difficulties (Whitlock 2008). For example, true Q_ST_ comparisons must obtain estimates of additive genetic variance, which means controlled crosses of dozens of pairs from each population. Doing so for six populations for our previous work was challenging, doing so for more than thirty populations would be prohibitive. Given the caveats that would be necessary with this dataset (i.e. having to use P_ST_ instead of Q_ST_), we decided not to pursue this analysis here.

### Population genetics

Our population genetic results indicate that there is very little geographic population structure in both continents. Population differentiation in neutral markers as measured by F_ST_ was low for both the native (0.0639) and invasive (0.0566) ranges. Past work on ragweed has found similarly low F_ST_ values (Genton *et al.* 2005; Chun *et al.* 2010; 2011; Martin 2011; Martin *et al.* 2016; van Boheemen *et al.* 2017). Martin and colleagues used SNPs to assess population differentiation of North American ragweed, and found that a solitary genetic cluster was the most likely population structure (Martin et al. 2016). Our STRUCTURE analyses showed the highest likelihoods for multiple clusters, but there was not a geographic pattern to the ancestral groupings, especially for the European populations. The consistent findings of low isolation by distance and population structure may not be surprising given that ragweed is a wind-pollinated outcrosser (Friedman and Barrett 2008).

The assumption that introduced species will always face significant deficits in genetic variation has been challenged by numerous counter-examples in the literature (Colautti & Lau 2015; Estoup *et al.* 2016). In some cases, due to multiple introductions and subsequent admixture, molecular diversity is actually higher in invasive populations when compared to their native counterparts (Novak & Mack 1993; Dlugosch & Parker 2008; Keller & Taylor 2008; 2010). Here, we estimated slightly higher metrics of genetic diversity (i.e., expected heterozygosity) in the invasive range for the represented populations. These findings corroborate a study in the French and North American ranges using microsatellite genetic variability where within population diversity was higher in the invasive range and overall genetic diversity was comparable between the two ranges (Genton *et al.* 2005). The high genetic diversity in the invasive range is likely the result of high propagule introduction from multiple sources in the native range (Genton et al. 2005a). The persistence of large, genetically diverse populations in the worst affected areas of Europe are dangerous sources for the spread of ragweed into currently unaffected areas. Roads and railway tracks are ideal corridors and habitats for ragweed and could facilitate multiple introductions and gene flow (Lavoie *et al.* 2007).

### Conclusions

Adaptation in introduced environments is not just theoretically interesting, but also has extremely important ecological and economic implications. Gradients in abiotic variables can lead to divergent selection across an introduced range and, if populations have sufficient genetic variation, create clines in traits (Maron *et al.* 2004). This adaptation can exacerbate the negative impacts of introduced species (Maron et al. 2004; Huey et al. 2005)

Ragweed has already caused economic and health impacts in Europe (Buttenschøn et al. 2010). Climate change is expected to extend its growing season, and it continues to spread its geographic range (Ziska et al. 2011). Our research corroborates past findings that ragweed has been able to adapt to its invasive range (Chun et al. 2011; Hodgins and Rieseberg 2011). The genetic, phenotypic and ecological traits of introduced ragweed make it very likely that the invasion will worsen.

## Acknowledgments

Bruce Hall, Bill Cole and Andrew Petrie provided excellent greenhouse support. Stephan Schneider provided much helpful assistance and encouragement at KSR. Antonio DiTomasso, Kathleen Howard, Rakesh Chandran, Quentin Martinez and Barbara Tokarska-Guzik helped to locate populations, assisted with collections and generously offered their very considerable expertise. Bruno Chauvel, Jannice Friedman, Peter Kotanen and all provided invaluable ragweed advice. Ali Parker, Amanda Gorton, Elisabeth McGregor, Rafal Janik and Valérie Kaakeh helped with collections. We are very grateful to all the authors of the tools that we used for this study, and help from many dedicated undergraduates. We thank Laura Galloway, Emily Josephs, Peter Kotanen, Art Weis, Stephen Wright for comments that improved this manuscript.

## Appendices

**Table A1.**
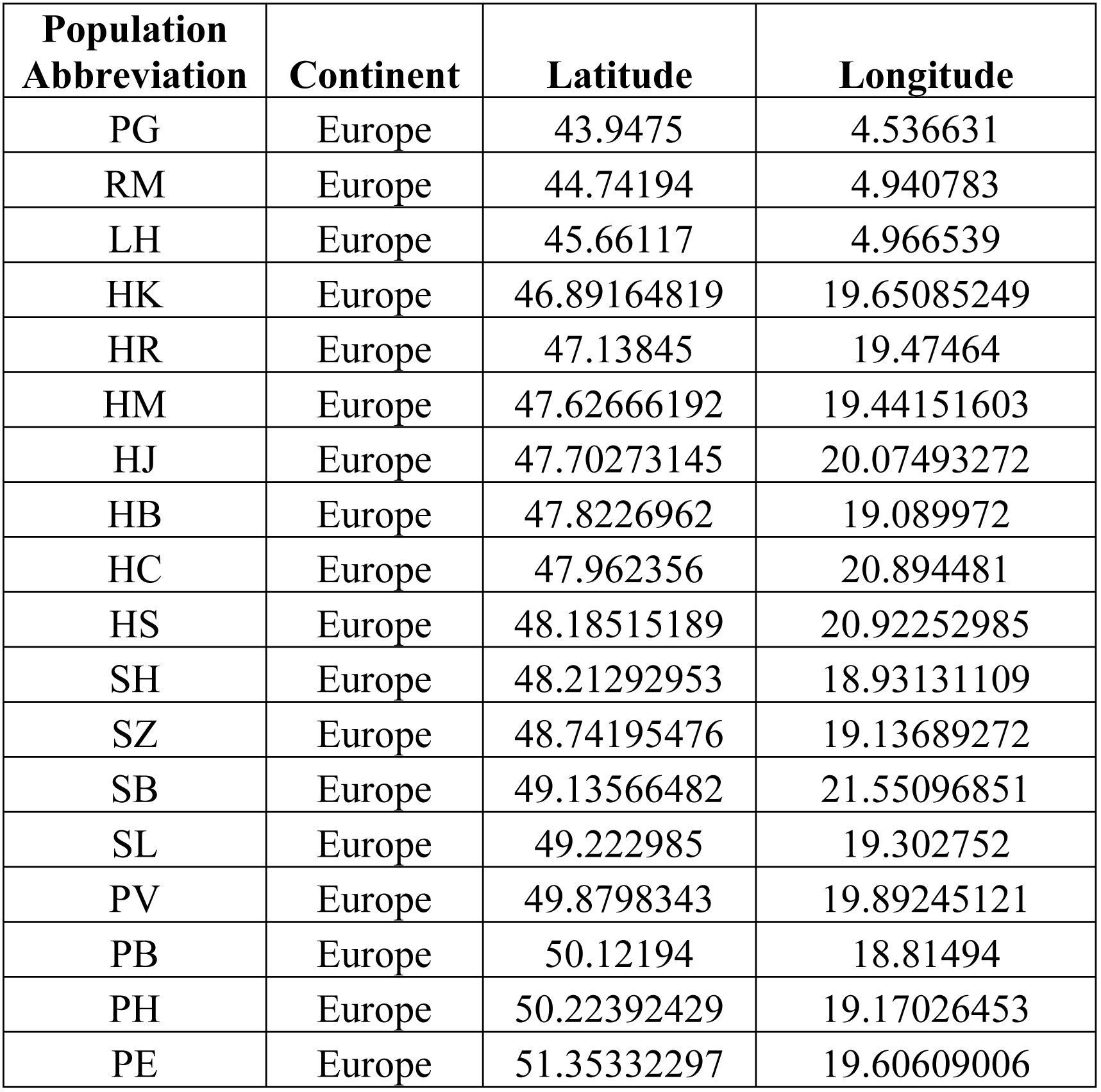
Population coordinates for 18 European (invasive) ragweed populations.

**Table A2.**
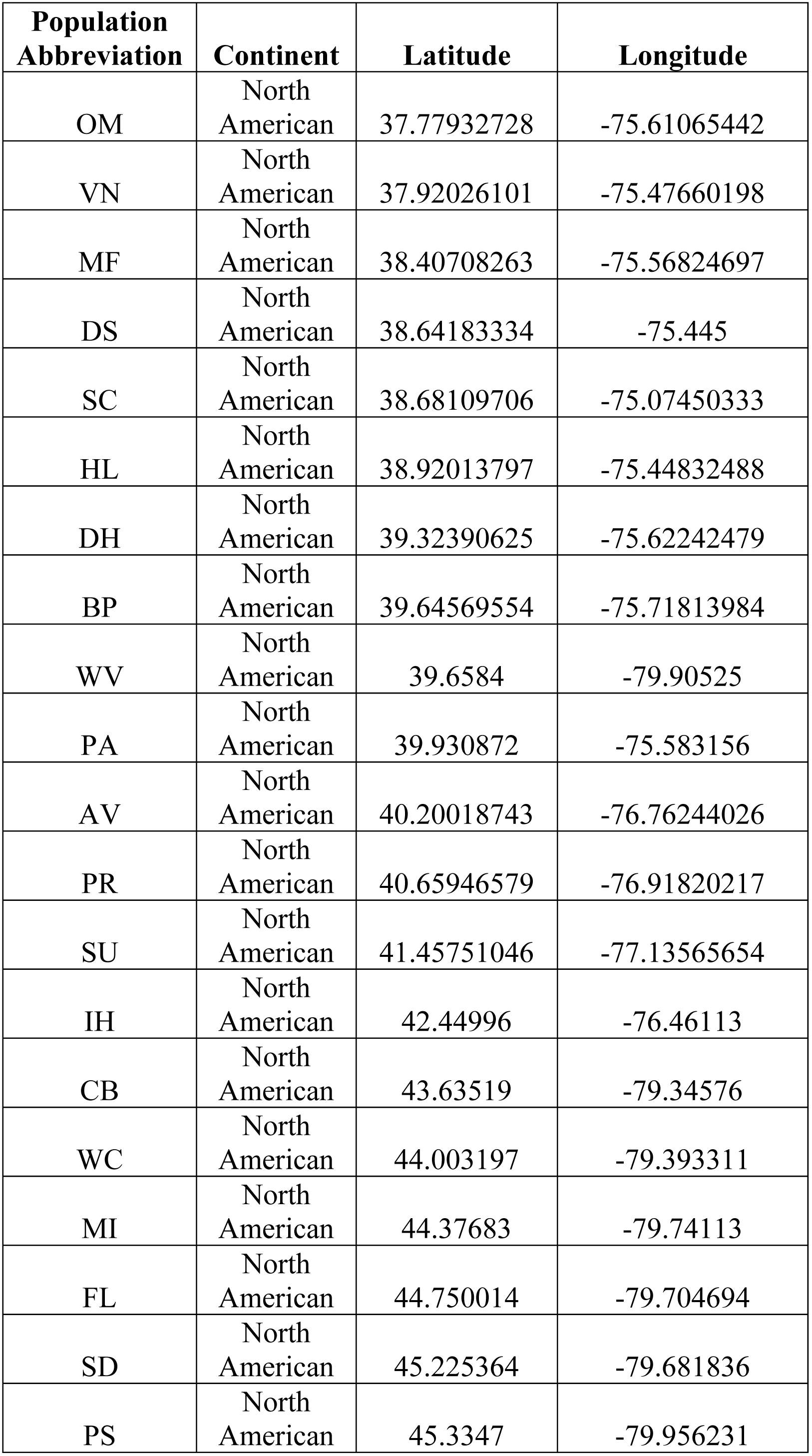
Population coordinates for 20 North American (native) ragweed populations.

**Table A3.** Pairwise F_st_ values for North American ragweed populations.

|  | VN | MF | HL | BP | WV | PA | AV | PR | SU | CB | MI | FL | SD | PS | DH | IH | WC |
| --- | --- | --- | --- | --- | --- | --- | --- | --- | --- | --- | --- | --- | --- | --- | --- | --- | --- |
| VN | -- | 0.061 | 0.101 | 0.18 | 0.078 | 0.082 | 0.18 | 0.089 | 0.093 | 0.075 | 0.073 | 0.061 | 0.13 | 0.085 | 0.086 | 0.066 | 0.063 |
| MF | 0.061 | -- | 0.033 | 0.118 | 0.02 | 0.018 | 0.116 | 0.031 | 0.029 | 0.017 | 0.018 | 0.022 | 0.049 | 0.027 | 0.02 | 0.007 | 0.015 |
| HL | 0.101 | 0.033 | -- | 0.154 | 0.051 | 0.052 | 0.149 | 0.059 | 0.061 | 0.049 | 0.05 | 0.054 | 0.092 | 0.057 | 0.053 | 0.04 | 0.047 |
| BP | 0.18 | 0.118 | 0.154 | -- | 0.131 | 0.131 | 0.213 | 0.137 | 0.145 | 0.129 | 0.139 | 0.13 | 0.188 | 0.145 | 0.139 | 0.117 | 0.122 |
| WV | 0.078 | 0.02 | 0.051 | 0.131 | -- | 0.034 | 0.125 | 0.044 | 0.038 | 0.024 | 0.019 | 0.026 | 0.049 | 0.03 | 0.043 | 0.02 | 0.02 |
| PA | 0.082 | 0.018 | 0.052 | 0.131 | 0.034 | -- | 0.125 | 0.043 | 0.043 | 0.033 | 0.032 | 0.035 | 0.068 | 0.044 | 0.042 | 0.021 | 0.029 |
| AV | 0.18 | 0.116 | 0.149 | 0.213 | 0.125 | 0.125 | -- | 0.133 | 0.135 | 0.124 | 0.129 | 0.121 | 0.171 | 0.137 | 0.136 | 0.112 | 0.114 |
| PR | 0.089 | 0.031 | 0.059 | 0.137 | 0.044 | 0.043 | 0.133 | -- | 0.051 | 0.04 | 0.039 | 0.042 | 0.072 | 0.049 | 0.053 | 0.031 | 0.036 |
| SU | 0.093 | 0.029 | 0.061 | 0.145 | 0.038 | 0.043 | 0.135 | 0.051 | -- | 0.035 | 0.036 | 0.036 | 0.079 | 0.044 | 0.05 | 0.025 | 0.031 |
| CB | 0.075 | 0.017 | 0.049 | 0.129 | 0.024 | 0.033 | 0.124 | 0.04 | 0.035 | -- | 0.009 | 0.009 | 0.046 | 0.02 | 0.043 | 0.015 | 0.007 |
| MI | 0.073 | 0.018 | 0.05 | 0.139 | 0.019 | 0.032 | 0.129 | 0.039 | 0.036 | 0.009 | -- | 0.01 | 0.041 | 0.011 | 0.042 | 0.016 | 0.003 |
| FL | 0.061 | 0.022 | 0.054 | 0.13 | 0.026 | 0.035 | 0.121 | 0.042 | 0.036 | 0.009 | 0.01 | -- | 0.04 | 0.017 | 0.041 | 0.017 | 0.006 |
| SD | 0.13 | 0.049 | 0.092 | 0.188 | 0.049 | 0.068 | 0.171 | 0.072 | 0.079 | 0.046 | 0.041 | 0.04 | -- | 0.054 | 0.071 | 0.047 | 0.036 |
| PS | 0.085 | 0.027 | 0.057 | 0.145 | 0.03 | 0.044 | 0.137 | 0.049 | 0.044 | 0.02 | 0.011 | 0.017 | 0.054 | -- | 0.048 | 0.025 | 0.012 |
| DH | 0.086 | 0.02 | 0.053 | 0.139 | 0.043 | 0.042 | 0.136 | 0.053 | 0.05 | 0.043 | 0.042 | 0.041 | 0.071 | 0.048 | -- | 0.031 | 0.037 |
| IH | 0.066 | 0.007 | 0.04 | 0.117 | 0.02 | 0.021 | 0.112 | 0.031 | 0.025 | 0.015 | 0.016 | 0.017 | 0.047 | 0.025 | 0.031 | -- | 0.009 |
| WC | 0.063 | 0.015 | 0.047 | 0.122 | 0.02 | 0.029 | 0.114 | 0.036 | 0.031 | 0.007 | 0.003 | 0.006 | 0.036 | 0.012 | 0.037 | 0.009 | -- |

**Table A4.** Pairwise F_st_ values for European ragweed populations.

|  | PG | LH | HK | HR | HJ | HB | HC | HS | SZ |
| --- | --- | --- | --- | --- | --- | --- | --- | --- | --- |
| PG | -- | 0.021 | 0.050 | 0.026 | 0.091 | 0.051 | 0.043 | 0.038 | 0.054 |
| LH | 0.021 | -- | 0.035 | 0.018 | 0.083 | 0.034 | 0.035 | 0.030 | 0.047 |
| HK | 0.050 | 0.035 | -- | 0.036 | 0.105 | 0.063 | 0.056 | 0.050 | 0.07 |
| HR | 0.026 | 0.018 | 0.036 | -- | 0.080 | 0.037 | 0.039 | 0.028 | 0.046 |
| HJ | 0.091 | 0.083 | 0.105 | 0.080 | -- | 0.120 | 0.099 | 0.092 | 0.114 |
| HB | 0.051 | 0.034 | 0.063 | 0.037 | 0.120 | -- | 0.061 | 0.051 | 0.071 |
| HC | 0.043 | 0.035 | 0.056 | 0.039 | 0.099 | 0.061 | -- | 0.046 | 0.063 |
| HS | 0.038 | 0.030 | 0.050 | 0.028 | 0.092 | 0.051 | 0.046 | -- | 0.058 |
| SZ | 0.054 | 0.047 | 0.070 | 0.046 | 0.114 | 0.071 | 0.063 | 0.058 | -- |

**Table A5.** Population genetic summary statistics for North American ragweed.

| Statistic | Value | Std.Dev. | c.i.2.5% | c.i.97.5% | Description |
| --- | --- | --- | --- | --- | --- |
| Num | 2 | 0 | 2 | 2 | Number of alleles |
| Eff_num | 1.163 | 0.001 | 1.161 | 1.165 | Effective number of alleles |
| Ho | 0.108 | 0.001 | 0.107 | 0.11 | Observed heterozygosity |
| Hs | 0.141 | 0.001 | 0.14 | 0.142 | Heterozygosity within populations |
| Ht | 0.15 | 0.001 | 0.148 | 0.151 | Total heterozygosity |
| H't | 0.15 | 0.001 | 0.149 | 0.152 | Corrected total heterozygosity |
| Gis | 0.23 | 0.001 | 0.228 | 0.233 | Inbreeding coefficient |
| Gst | 0.061 | 0.001 | 0.06 | 0.062 | Fixation index |
| G'st(Nei) | 0.064 | 0.001 | 0.063 | 0.065 | Nei, corrected fixation index |
| G'st(Hed) | 0.071 | 0.001 | 0.07 | 0.072 | Hedrick, standardised fixation index |
| G"st | 0.075 | 0.001 | 0.073 | 0.076 | Corrected standardised fixation index |
| D_est | 0.011 | 0 | 0.011 | 0.011 | Jost, differentiation |

**Table A6.** Population genetic summary statistics for European ragweed.

| Statistic | Value | Std.Dev. | c.i.2.5% | c.i.97.5% | Description |
| --- | --- | --- | --- | --- | --- |
| Num | 2 | 0 | 2 | 2 | Number of alleles |
| Eff_num | 1.224 | 0.001 | 1.221 | 1.226 | Effective number of alleles |
| Ho | 0.137 | 0.001 | 0.136 | 0.138 | Observed heterozygosity |
| Hs | 0.192 | 0.001 | 0.19 | 0.193 | Heterozygosity within populations |
| Ht | 0.202 | 0.001 | 0.2 | 0.204 | Total heterozygosity |
| H't | 0.203 | 0.001 | 0.201 | 0.205 | Corrected total heterozygosity |
| Gis | 0.285 | 0.002 | 0.281 | 0.289 | Inbreeding coefficient |
| Gst | 0.051 | 0.001 | 0.05 | 0.052 | Fixation index |
| G'st(Nei) | 0.057 | 0.001 | 0.055 | 0.059 | Nei, corrected fixation index |
| G'st(Hed) | 0.065 | 0.001 | 0.063 | 0.066 | Hedrick, standardised fixation index |
| G"st | 0.071 | 0.001 | 0.069 | 0.073 | Corrected standardised fixation index |
| D_est | 0.014 | 0 | 0.014 | 0.015 | Jost, differentiation |

**Table A7.** AMOVA output for North American and European ragweed.

| Source of Variation | Nested in | %var | F-stat | F-value | Std.Dev. | c.i.2.5% | c.i.97.5% | P-value | F'-value |
| --- | --- | --- | --- | --- | --- | --- | --- | --- | --- |
| Within Individual | -- | 0.795 | F<sub>it</sub> | 0.205 | 0.001 | 0.204 | 0.207 | -- | -- |
| Among Individual | Population | 0.162 | F<sub>is</sub> | 0.169 | 0.001 | 0.167 | 0.171 | 0.001 | -- |
| Among Population | Continent | 0.042 | F<sub>sc</sub> | 0.042 | 0 | 0.041 | 0.043 | 0.001 | 0.048 |
| Among Continents | -- | 0.002 | F<sub>ct</sub> | 0.002 | 0 | 0.001 | 0.002 | 0.017 | 0.002 |

**Figure A1.**
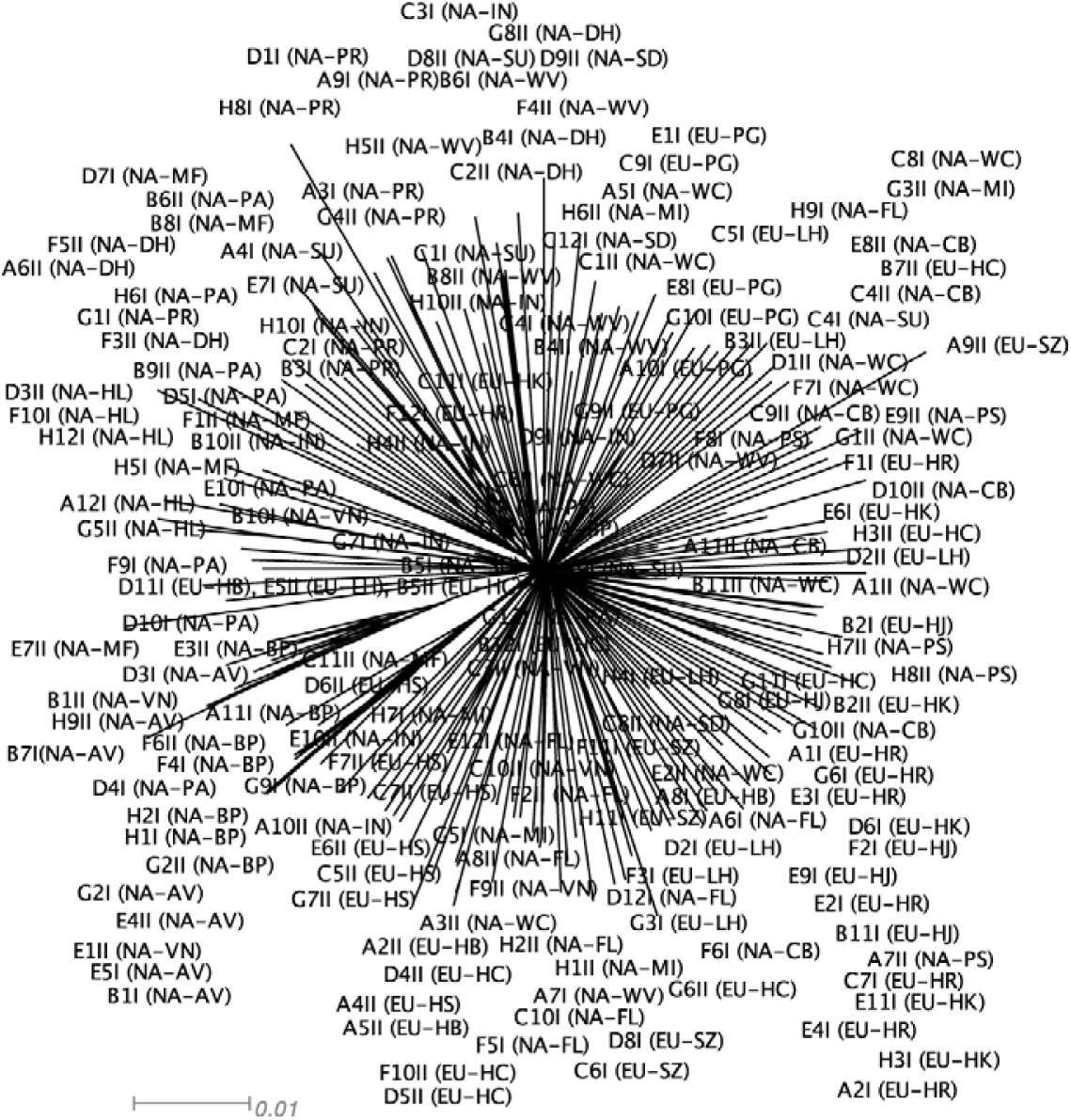
Splitstree results for invasive and native ragweed populations.

**Figure A2.**
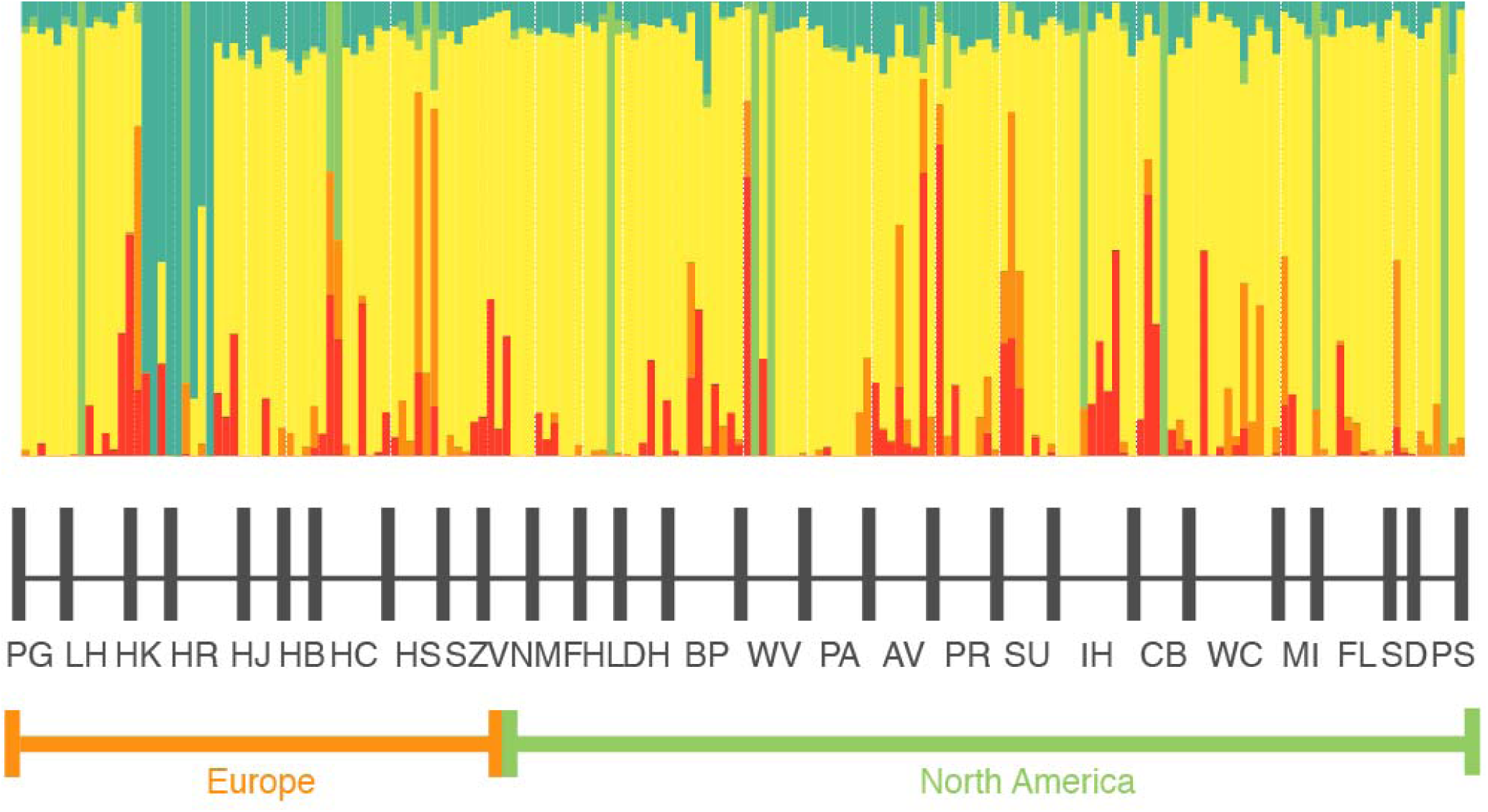
STRUCTURE results for all ragweed populations. Populations are ordered first by continent and then from lowest to highest latitudes within continents.

